# Predictive remapping of visual features beyond saccadic targets

**DOI:** 10.1101/297481

**Authors:** Tao He, Matthias Fritsche, Floris P. de Lange

## Abstract

Visual stability is thought to be mediated by predictive remapping of the relevant object information from its current, pre-saccadic locations to its future, post-saccadic location on the retina. However, it is heavily debated whether and what feature information is predictively remapped during the pre-saccadic interval. Using an orientation adaptation paradigm, we investigated whether predictive remapping occurs for stimulus features and whether adaptation itself is remapped. We found strong evidence for predictive remapping of a stimulus presented shortly before saccade onset, but no remapping of adaptation. Furthermore, we establish that predictive remapping also occurs for stimuli that are not saccade targets, pointing toward a ‘forward remapping’ process operating across the whole visual field. Together, our findings suggest that predictive feature remapping of object information plays an important role in mediating visual stability.

## Introduction

Each time we move our eyes, the image of objects in the world shifts its position on the retina, yet our perception is remarkably stable. Previous research revealed that predictive remapping could contribute to this visual stability. Predictive remapping refers to the phenomenon that neurons become active in response to stimuli outside their receptive fields (RFs) shortly before a saccade moves their receptive fields onto the stimulated regions (Duhamel et al., 1992). Predictive remapping has been demonstrated in many cortical regions, such as the lateral intraparietal area (LIP) (Duhamel et al., 1992), the frontal eye field (FEF) (Goldberg and Bruce, 1990; Umeno and Goldberg, 1997), superior colliculus (SC) (Walker et al., 1995), and early visual cortex including V2, V3 and V3a (Nakamura and Colby, 2002). Predictively increasing activity of visually responsive neurons in these areas according to postsaccadic stimulus information could facilitate the processing of visual information across saccades, which is crucial for achieving perceptual stability.

Although predictive remapping has been widely studied, there is an ongoing debate regarding the issue whether and how feature information of visual objects is remapped during this process (Cavanagh et al., 2010; Ezzati et al., 2008; Harrison et al., 2013; He et al., 2017; Lescroart et al., 2016; Mayo and Sommer, 2010; Melcher, 2010, 2007, 2005; Pelli and Cavanagh, 2013; Zimmermann et al., 2017; Zirnsak and Moore, 2014). On the one hand, several psychophysical studies suggest that visual feature information, such as orientation and letter information, is transmitted around the time of a saccade (Harrison et al., 2013; He et al., 2017; Melcher, 2007). More specifically, it is suggested that relevant features of a test stimulus, which are extracted before the saccade, are transferred to their postsaccadic retinal location based on the computation of the saccade vector. On the other hand, Rolfs et al., (2011) proposed that it is merely the attentional pointers, but not the feature information, that are predictively remapped across saccades. By linking the attentional pointers at the current and future retinotopic locations together, the feature information at these two distinct locations is combined at higher processing stages.

The tilt aftereffect (TAE), in which prolonged exposure to a stimulus (the adaptor) results in a perceptual shift of a test stimulus away from the adaptor is a sensitive method to address the question of feature remapping (Knapen et al., 2011; Melcher, 2007). Namely, orientation feature integration between the pre- and post-saccadic location can be inferred from observing a TAE. There has been considerable confusion however concerning *what* is supposedly remapped prior to executing a saccade. Specifically, it is unclear whether the adaptor (or the state of adaptation), the test stimulus, or both, is remapped (see Figure 1C and 1D). Moreover, the spatial properties of remapping are a current topic of debate. In particular, it is not clear whether receptive fields are shifted to their postsaccadic location, or toward the saccade target (Zirnsak et al., 2014; Zirnsak and Moore, 2014). Since in most of the previous studies the probe location coincided with the saccade target location, these previous studies are unable to differentiate between convergent and forward remapping effects (Marino and Mazer, 2016; Neupane et al., 2016a), and studies that aimed to dissociate these effects provided conflicting results (Neupane et al., 2016b; Zirnsak et al., 2014).

**Figure 1.**
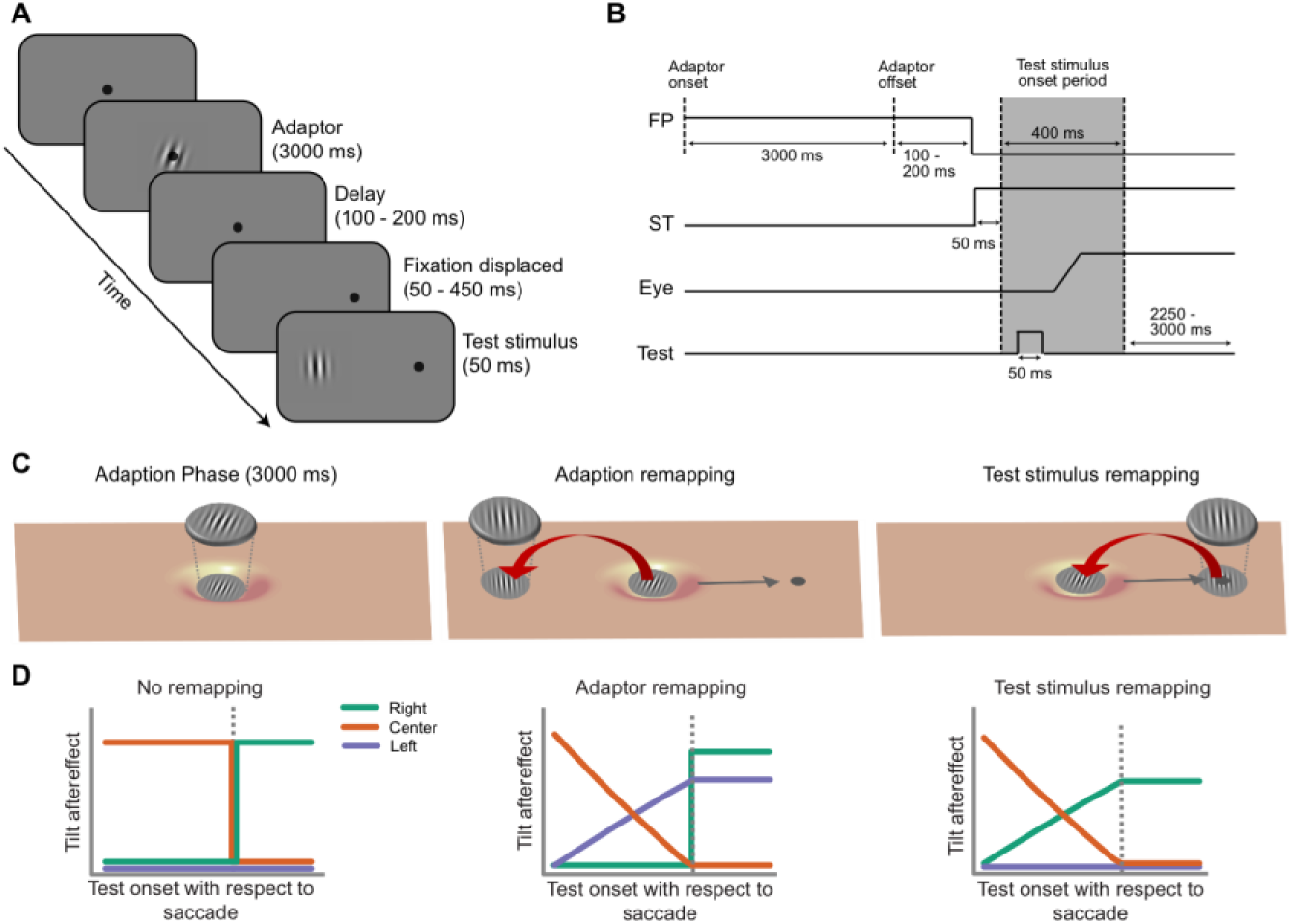
Experimental design and hypotheses. **A)** Experimental design. An adaptor was first presented at the initial fixation location for 3 s. After a random delay period, participants were asked to make a horizontal eye movement to the saccade target following the shift of fixation (black dot). Immediately after the presentation of test stimulus, participants were asked to report whether the test stimulus was tilted to the left or right relative to vertical. The test stimulus could appear at one of three locations (left, center, or right) and could appear before or after saccade onset. **B)** Time course of a trial. The grey area is the potential time period that a test stimulus could be presented (before, during or after a saccade). Trials on which a test stimulus was presented during the saccade were removed prior to the analysis. FP, fixation point; ST, saccade target. **C)** Left: Adaptation effect. After a prolonged exposure to the adaptor, the neural population that is sensitive to the location of the adaptor becomes adapted (the “hole” in the figure). Middle: Adaptator remapping hypothesis. Upon preparing a rightward saccade (grey arrow), the adaptator is remapped to its anticipated postsaccadic location. As a consequence, a TAE is expected at the left location. Right: Test stimulus remapping hypothesis. Upon preparing a rightward saccade (grey arrow), the test stimulus is remapped to its anticipated postsaccadic location. As a consequence, a TAE is expected at the right location. **D)** The expected pattern of results under conditions of no remapping (left column), adaptor remapping (middle column) and test stimulus remapping (right column).

In the current study we investigated whether stimulus orientation is predictively remapped, and whether adaptation itself is remapped, as has been suggested before. Further, we examined whether presaccadic remapping also occurs for non-saccade targets, in order to distinguish between forward and convergent remapping. To this end, we made use of the orientation adaptation paradigm to test the TAE at each critical location, e.g., the initial fixation location, the saccade target location and the remapped location of adaptor. To preview, we found that predictive feature remapping only occurs for stimuli presented shortly before a saccade and irrespective of whether they are a saccade goal, suggesting that the visual system employs forward predictive feature remapping to mediate visual stability.

## Results

We collected psychophysical data in a series of three experiments, each employing 24 human participants. In total, we recruited a sample of 72 participants and 82,080 trials.

### Selective remapping of future target stimuli but not adaptation

Our first aim was to test whether the test stimulus or adaptation is remapped. To this end, we used a modified version of an adaptation paradigm that has been previously described (Melcher, 2007), while monitoring eye position (Figure 1A). During the experiment (Experiment 1), participants maintained fixation in the central screen while they adapted to an oriented Gabor (+20° or -20°). When the central fixation was displaced to a peripheral location, the participants quickly moved their eyes to track the dot. The test stimulus (Gabor stimulus with one of five orientations: -2°, -1°, 0°, 1°, 2°) could be presented at one of three potential locations (i.e., the initial fixation location, the saccade target location or the remapped location of adaptor). Participants indicated whether it was rotated clockwise or counterclockwise relative to vertical by pressing one of two buttons.

When a test stimulus was presented well before the saccade initiation (Figure 2, left column), we found that the perceived orientation of the test stimulus at each location was systematically biased away from the adaptor stimulus that was previously presented at the center of the screen (Figure 2). This repulsive bias, which is well known as the tilt-aftereffect (TAE) in orientation perception, was quantified as the difference in the point of subjective equality (PSE) between a left-tilted and right tilted adaptor (illustrated as the black bar between the psychometric curves). It was strongest when the test stimulus was presented at the center location (middle row), where the adaptor had been presented, and markedly reduced but still present at the other two locations. We next investigated if, when, and where the TAE was transferred shortly before subjects initiated a saccade. We found that shortly before an eye movement, the TAE was significantly reduced at the remapped location of the adaptor (Figure 2, “Left” location, violet lines, comparison between first and second time point: *p* = 0.0165). Also at the initial fixation location, the TAE was reduced before an eye movement (Figure 2, “Center” location, orange lines, comparison between first and second time point: *p* = 0.0039). However, the TAE at the future saccade target location was significantly enhanced before the onset of the saccade (Figure 2, “Right” location, green lines, comparison between first and second time point: *p* < 0.0001). This opposite behavior over time between the locations resulted in a significant (*p* = 0.0002) interaction between target location (initial vs. future saccade location) and time (first vs. second time bin), showing that TAE increased at the future saccade target location and decreased at the initial location.

**Figure 2.**
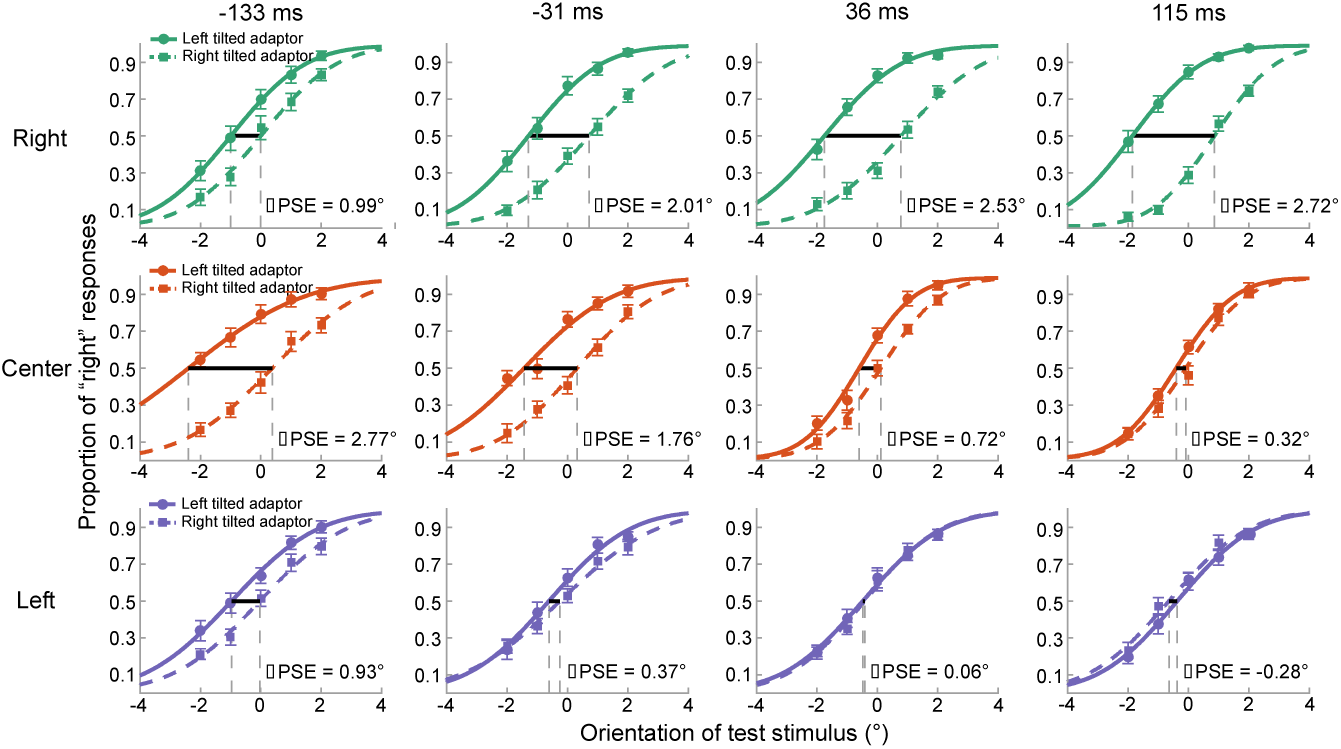
Psychometric curves for orientation judgements in Experiment 1. The number above each column represents the mean test stimulus onset relative to saccade onset (first and second columns) or offset (third and fourth columns) for each time bin. The labels of “Left”, “Center” and “Right” in the left of each row represent the three different test stimulus locations. For each panel, we plotted the percentage of a “right” response (y axis) as a function of the orientation of a test stimulus (x axis) for each time bin and location. the positive x values mean the test stimulus was tilted more clockwise while the negative x values mean more countclockwise relative to vertical. The black lines indicate that the difference between the point of subjective equality (PSE, the angle in which participants judge a test stimulus was oriented left or right equally) of the leftward (solid line) and rightward (dashed line) tilted adaptor conditions, the value of tilt aftereffect (TAE) was defined as the half of ∆PSE.

These results are consistent with, and extend, those reported by Melcher (2007). When the test stimulus was presented at the saccadic target location, the features of the test stimulus were predictively remapped to the presaccadic foveal location, that was previously adapted. Importantly, however, we found no TAE at the location where the adaptor would hypothetically be remapped. Put simply, it is the orientation feature information of a stimulus that is perceived shortly before the saccade, but not a previously seen adaptor and its consequences, that is predictively remapped before saccade onset.

### Selective remapping of non-saccade targets

In Experiment 1, we observed predictive feature remapping of the test stimulus towards its post-saccadic location. However, since in this crucial condition the test stimulus was always a saccade target, we cannot differentiate between a mechanism that only remaps saccade targets to the fovea (convergent remapping) and one that more generally remaps stimuli across the visual field to their post saccadic locations (forward remapping). In order to test whether remapping also occurs for non-saccade targets, we flashed both the adaptor and test stimulus 4° vertically above fixation. The idea behind this design is straightforward: If predictive remapping only occurs for saccade targets, we would expect no TAE when the test stimulus is presented 4° above the fixation target. However, if predictive remapping also occurs for stimuli that are not saccade targets, an increase of TAE for peripherally presented test stimuli should be observable during the pre-saccadic period.

Despite the fact that different locations were used for the adaptor and the test stimulus, we found a similar pattern of results as in Experiment 1. Specifically, before an eye movement, the TAE was significantly increased at the future saccade target location (Figure 3, “Right” location, green lines, comparison between first and second time point: *p* = 0.0068). However, the TAE at the initial fixation location was decreased for the second compared to the first time bin (Figure 3, “Center” location, orange lines, comparison between first and second time point: *p* = 0.0197). This opposite behavior over time between the locations also resulted in a significant (*p* = 0.0019) interaction between test stimulus location (initial vs. future fixation location) and time (first vs. second time bin). This result suggests that predictive remapping likewise occurs for stimuli that are not saccade targets, consistent with a forward remapping account.

**Figure 3.**
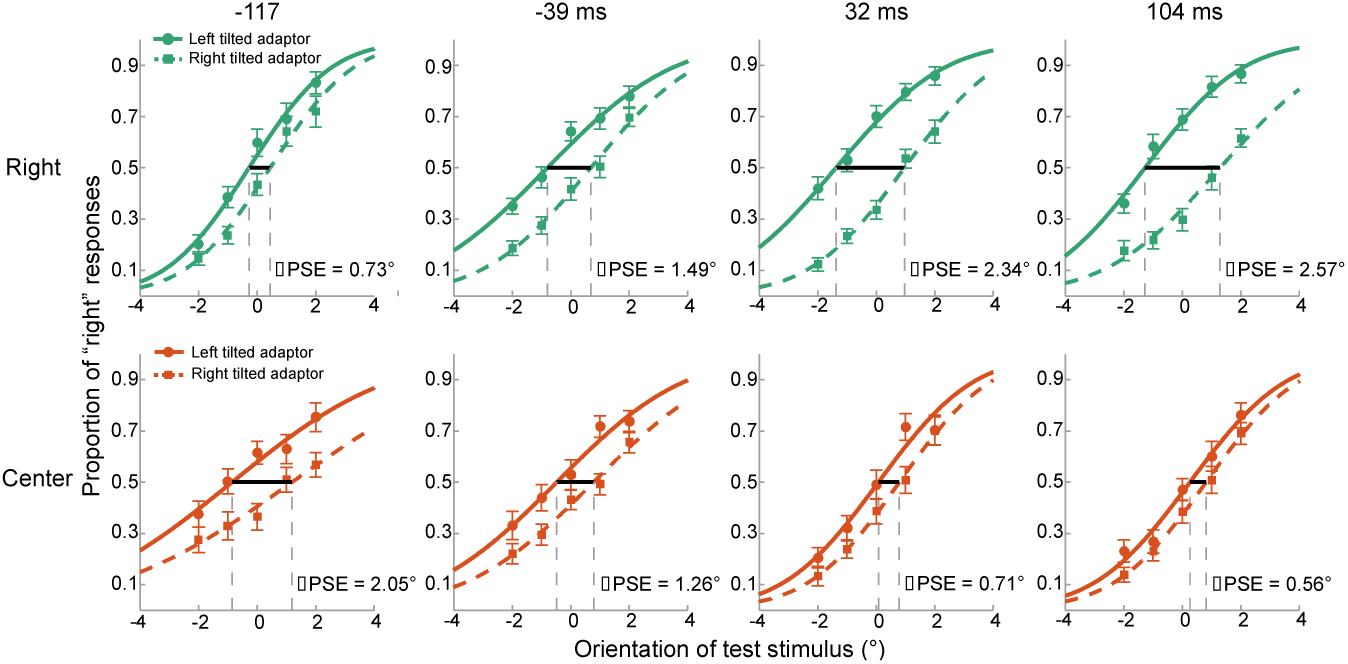
Psychometric curves for orientation judgements in Experiment 2. Same conventions as in Figure 2 but only “Right” and “Center” location were tested, while both the adaptor and test stimulus were presented 4° above fixation.

The results of Experiment 1 and 2 indicate that predictive remapping of orientation occurs, irrespective of whether the stimulus is a saccade target. However, due to the short spatial distance between the test stimulus and saccade target location, and between the adaptor location and foveal fixation (both 4°), one may still argue that the findings in Experiment 2 could be explained by remapping of stimuli close to the fixation target or the fovea. To more directly contrast the convergent remapping and forward remapping hypotheses, we designed Experiment 3, in which two oppositely oriented adaptors were presented simultaneously at peripheral locations while the test stimulus was flashed at the initial fixation location or saccade target location.

Forward remapping hypothesizes a remapping of receptive fields in the same direction as the saccade. Therefore, a test stimulus presented at the initial fixation location (center) will, just prior to making a rightward saccade, be remapped to the left, opposite of the saccade vector. Convergent remapping, on the other hand, hypothesizes a remapping of receptive fields towards the future saccade location. In this case, no TAE would be predicted for a stimulus presented at the initial fixation location, because no receptive fields are remapped to this location (see Materials and Methods for more details).

The results indicated a positive TAE for stimuli presented at the “Center” location, just before participants made a saccade (Figure 4, orange lines, second column: *p* = 0.0337), in line with forward remapping. Conversely, the TAE for stimuli presented at the saccade target location (“Right” location) was significantly decreased before saccade onset (Figure 4, green lines, comparison between first and second time point: *p* < 0.0001). This pattern of results suggests that the test stimulus was predictively forward remapped prior to the eye movement.

**Figure 4.**
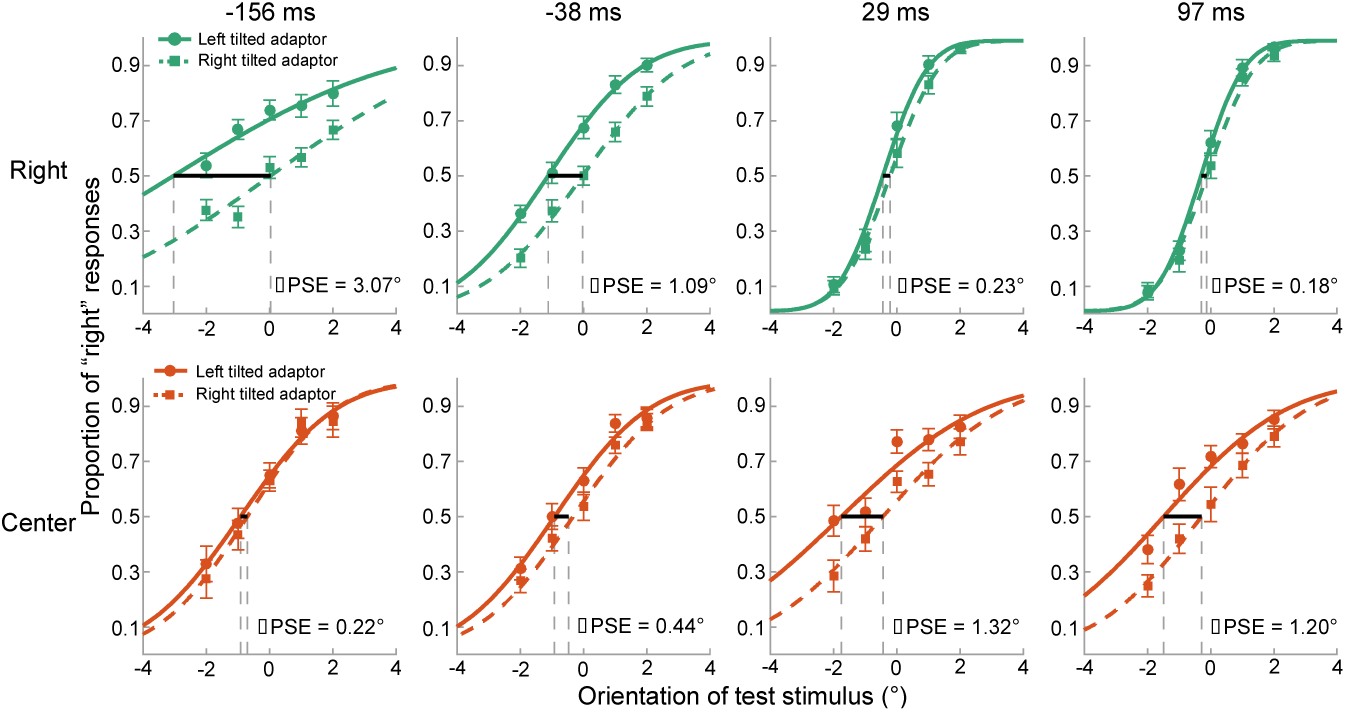
Psychometric curves for orientation judgements in Experiment 3. Same conventions as in Figure 2 but only “Right” and “Center” location were tested while two oppositely oriented adaptors were presented in the periphery before saccade target onset.

## Discussion

We used an orientation adaptation paradigm to investigate whether and how feature information is predictively remapped prior to saccades. In Experiment 1 (see Figure 5A), we found strong evidence for predictive remapping of visual information that is presented shortly before saccade onset, but no remapping of adaptation, as had been previously hypothesized (Melcher, 2007; Rolfs et al., 2011). In Experiment 2 and 3 (see Figure 5B and Figure 5C), we provided evidence that pre-saccadic remapping of features also occurs for stimuli that are not a saccade target, consistent with forward remapping, which further underscores the generality of this mechanism (Neupane et al., 2016a, 2016b).

**Figure 5.**
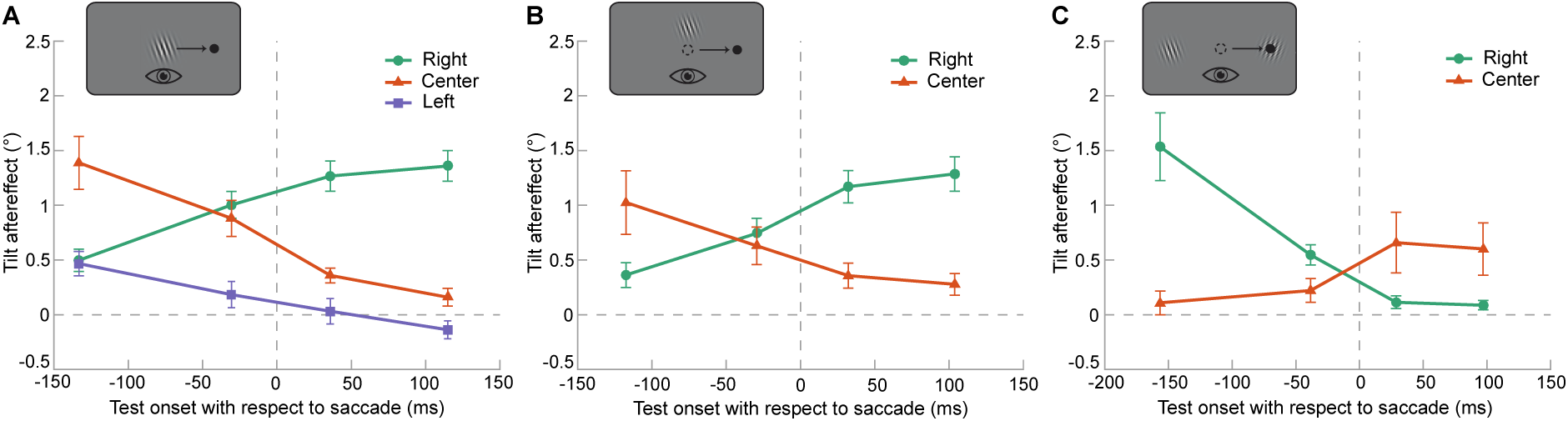
Comparison of Tilt aftereffect (TAE) across experiments. **A)** Experiment 1: the adaptor was shown at fixation location. For analysis, leftward saccade trials were collapsed into rightward saccade trials. The vertical dashed line indicates the onset/offset of a saccade. Error bar represent one SD of the bootstrapped distribution. **B)** Experiment 2: the adaptor was shown 4° above fixation. Only initial fixation and saccade target location were probed. Other parameters were identical to Experiment 1. **C)** Experiment 3: two oppositely oriented adaptors were presented at peripheral location. The test stimulus was presented at the initial fixation location or saccade target location. The error bar represents one SD of the bootstrapped distribution.

### No remapping of adaptation

The results of Experiment 1 and 3 indicate that while features of stimuli presented shortly before the impending saccade are remapped to their future retinal location, the adaptation effect itself is not remapped. In our experiments, the adaptor stimulus is presented during an initial fixation period, long before participants are instructed to prepare a saccade. Therefore, at the time participants could prepare a specific saccade plan, the adaptor stimulus had already disappeared. Since the saccade preparation occurred after the adaptor stimulus offset, any processing of the adaptor stimulus is likely finished by the time participants prepare the saccade. As the remapping dynamics also clearly show, only stimulus information that is presented very shortly before the saccade is remapped. This is also in line with the notion that adaptation occurs in a retinotopic reference frame (Knapen et al., 2011; Wenderoth and Wiese, 2008), possibly due to a reduction of excitability of the adapted neurons. It is unlikely that such a reduction of neuronal excitability can be ‘remapped’ by the planning of a saccade.

Contrary to our results, a recent paper by He et al. (2017) did observe feature remapping of adaptation. In their study, however, participants were required to make the same saccade on every trial, while the test stimulus also always appeared at the same location (i.e., the hypothetically remapped location of the adaptor). Therefore, it is highly likely that participants strongly attended this location, leading to an apparent remapping of the adaptation.

### Remapping of features or attentional pointers?

The question whether feature information is involved in the predictive remapping process has been extensively debated in the recent decade. Rolfs et al. (2011) found that visual performance was gradually enhanced at the future retinotopic location even before the onset of eye movements. Since the target was very difficult to detect and required a high degree of attention toward the particular location, the authors proposed that attention, rather than feature information is predictively remapped prior to a saccade. This hypothesis was further supported by several subsequent studies (Harrison et al., 2013; e.g., Hunt and Cavanagh, 2011; Jonikaitis et al., 2013; Puntiroli et al., 2015). However, in recent years a number of studies provide evidence that feature information, in addition to the attentional pointers alone, is also involved in transsaccadic remapping (Harrison and Bex, 2014; He et al., 2017; Zimmermann et al., 2017, 2016). Our study is in line with these studies, and further extends the findings by showing that orientation features of an actively processed stimulus, rather than the adaptation effects due to previous stimulation, are remapped.

Notably, several fMRI studies also shown evidence for predictive feature remapping. Zimmermann et al., (2016) found that visual feature information was dynamically remapped from a retinotopic coordinate into a spatiotopic coordinate system in ventral visual areas V3, V4, and VO. (Merriam et al., 2007) found remapping of information associated with the execution of eye movements not only in higher-order extrastriate areas (areas V3A, hV4) but also in V1 and V2, although smaller in magnitude, consistent with an earlier study in non-human primates (Nakamura and Colby, 2002). How is this feature information transferred within the visual system? A possible explanation for this might be that feature remapping is the effect of the combination of corollary discharge and bottom-up information. Activity elicited by the test stimulus could be remapped under the guidance of corollary discharge signals (Rao et al., 2016; Sommer and Wurtz, 2006; Sperry, 1950). The basic idea of corollary discharge is that when the motor system generates a movement command for muscles to produce a movement, a copy or corollary of this command will also be sent to other regions of the brain to inform them about the impending movement. Thus when a saccade is prepared by the oculomotor system, a corollary discharge signal containing information about the onset and target location of the imminent eye movement could be used to redirect the flow of feature information in visual cortex (Fries, 1984; Tolias et al., 2001). In particular, while the neurons whose receptive field cover the stimulus location will be activated by the bottom-up signal at first, this signal will be combined with the corollary discharge in extrastriate cortex and then, via the SC to neurons whose receptive field will overlap with the stimulus region after the eye movement.

### Convergent and forward predictive remapping

In their seminal study, Duhamel et al., (1992) reported that a set of LIP neurons predictively shift their receptive fields from their current location to their future retinotopic location prior to a saccade. This type of predictive remapping was termed *forward remapping*, since RF locations are shifted parallel to the saccade vector, as has been observed in several studies (Walker et al., 1995; Umeno and Goldberg, 2001; Nakamura and Colby, 2002). However, another type of predictive remapping has been proposed, which is termed *convergent remapping*, suggesting that the receptive fields shift toward the saccade target location rather than their postsaccadic location (Tolias et al., 2001; Zirnsak et al., 2014). Due to limitations in the experimental paradigms, forward and convergent remapping are sometimes difficult to distinguish. In particular, in many previous studies the test stimulus often constituted the saccade target, and in this case forward and convergent remapping theories make indistinguishable predictions.

In our current study, when the test stimulus was presented outside the saccade target location (Experiment 2 and 3), we still observed a robust forward pre-saccadic remapping effect. This result is in line with a previous electrophysiological study in V4 (Neupane et al., 2016b). In contrast, convergent remapping has been reported in FEF (Zirnsak et al., 2014). We speculate that the convergent remapping in FEF, which is a non-visual area, may not be functionally related to shifting of receptive fields but rather in anticipating and selecting relevant stimuli near the saccade target location, to facilitate processing of saccade targets. Conversely, for the visual system, maintaining stable representations of features across saccades is critical for seamless visually guided behaviors, which may be enabled by forward remapping.

## Conclusion

We found strong support for predictive remapping of the orientation feature of a test stimulus that was presented shortly before saccade onset. This pre-saccadic remapping also occurred for stimuli that were not saccade targets, and had the characteristics of a ‘forward remapping’ process that operates across the whole visual field. Thereby, forward predictive feature remapping may constitute an important mechanism for mediating visual stability.

## Materials and Methods

The current study consisted of three experiments. In the first experiment we tested whether predictive feature remapping occurs for stimuli that are saccade targets. In the second experiment we tested whether predictive feature remapping similarly occurs for peripheral stimuli that are not saccade targets. The third experiment acted as a control experiment to further corroborate the results of Experiment 1 and 2.

### Participants

A total of 72 subjects participated in three experiments, engaging in a total of 82,080 trials. Each experiment had 24 subjects (Exp.1: 11 females, mean age 23.6 years, range from 19 to 43 years; Exp.2: 16 females, mean age 22.8 years, range from 18 to 30 years; Exp.3: 15 females, mean age 24.4 years, range from 20 to 34 years). The sample size was based on an a priori power calculation, computing the required sample size to achieve a power of 0.80 to detect an effect size of Cohen’s d ≥ 0.6, at alpha = 0.05 for a within-subject comparison. All participants reported normal or corrected-to-normal vision, and were naive with respect to the purposes of the study. Participants were recruited from the institute’s subject pool in exchange for either monetary compensation or study credits. The experiments were approved by the Radboud University Institutional Review Board and were carried out in accordance with the guidelines expressed in the Declaration of Helsinki. Written informed consent was obtained from all participants prior to the start of the study.

### Apparatus

All stimuli were generated with custom scripts written in python and were presented on a 24-inch flat panel display (BenQ XL2420T, resolution 1920 × 1080, refresh rate: 60Hz). The visible area of the display measured 48° × 27° visual degrees at a viewing distance of about 64 cm. The participants’ head position was stabilized with a chin rest. Eye movements were monitored by an Eyelink 1000 plus (SR Research®) eye-tracker, sampling at 1000 Hz. Only the right eye was recorded. Saccade initiation was detected online, with a velocity threshold of 30°/s and an acceleration threshold of 8000°/s^2^. A 9-point calibration and validation procedure was conducted at the beginning of each block.

### Stimuli and Experimental Design

#### Experiment 1

Participants were tested in a quiet and dimly lit laboratory. Each trial began with the presentation of a fixation dot at the center of the screen. This fixation dot also served as the drift-correction target and remained visible until the participant’s gaze was within 1 visual degrees of it and the space bar was pressed. The sequence of events and time course in a single trial is illustrated in Figure 1A and 1B.

After the initiation of the trial a black fixation dot (diameter = 0.4°) and an oriented Gabor patch (oriented +20° or -20° relative to vertical) were presented at the center of the screen against a uniform mid-gray background for 3 seconds. The Gabor patch consisted of a sinusoidal wave grating (spatial frequency = 2 cycles/°; phase = 0.25; contrast = 1.0), windowed by a Gaussian envelope (SD 1.67°).

Participants were asked to fixate the dot until it disappeared. After 3 seconds, the Gabor patch disappeared and participants continued maintaining fixation at the central dot for a 100 – 200 ms delay. After the delay, the fixation dot was horizontally displaced to the left or right side of the screen (8 visual degrees), which served as a cue for participant to make a saccade to the new fixation location. A test stimulus (Gabor stimulus with one of five orientations: -2°, -1°, 0°, 1°, 2°) was then flashed briefly at one of three locations (left, center or right) for 50 ms. Crucially, the onset of the test stimulus varied in the range of 50 – 350 ms after the displacement of the fixation dot, such that it could occur either before or after the onset of the saccade, given that human saccade latency is estimated to lie around 200 ms (Robinson, 1964). The participant’s task was to indicate whether the test stimulus was tilted to left or right with respect to vertical, regardless of its location.

Participants completed 3 sessions of the task, comprising a total of 1260 trials. There were 210 trials for each combination of the two adaptor tilt orientations and three test stimulus locations. If the participant’s gaze deviated more than 2° from the central fixation dot during the adaption period, or landed at a location that was more than 2° away from the saccade target, auditory and visual feedback was given and the trial was aborted. All aborted trials were discarded and retested in a random order, until all trials were completed successfully.

#### Experiment 2

In order to test whether predictive feature remapping also occurs for stimuli that are not saccade targets, we repeated Experiment 1, but presented both adaptor and test stimulus 4° above fixation. Consequently, the test stimulus was never a saccade target. In addition, as Experiment 1 yielded no evidence of adaptor remapping prior to a saccade, we did not test for remapping at the location where the adaptor would hypothetically be remapped in this experiment.

The trial sequence in Experiment 2 was identical to Experiment 1. Each trial began with the presentation of an oriented Gabor patch 4° above central fixation. Participants were next asked to move their eyes to the periphery following the shift of the fixation dot. The test stimulus was flashed 4° above the initial fixation location or the saccade target location to measure transfer of feature information between these two locations. Experiment 2 consisted of 2 sessions. For each combination of the two test stimulus locations and adaptor tilt orientations 270 trials were collected, resulting in a total of 1080 trials.

#### Experiment 3

In Experiment 3, the task was similar to Experiment 1, except that two oppositely oriented adaptors were presented simultaneously at the two peripheral locations. In a given trial, participants initially fixated at the center of the screen, while two oppositely oriented Gabor patches (either +20°/-20° or -20°/+20° from vertical) were presented simultaneously for 3 seconds, 8° left and right of the center of the screen. Next participants were prompted to move their eyes to the left or right peripheral location, following the shift of the fixation dot. The test stimulus was either flashed at the initial fixation location or at the saccade target location. Experiment 3 consisted of 2 sessions. For each combination of the two test stimulus locations and adaptor tilt orientations 270 trials were collected, resulting in a total of 1080 trials.

The logic behind Experiment 3 is as follows. Imagine a trial in which the participant performs a saccade to the right peripheral location. Under the forward remapping hypothesis, a test stimulus that is centrally presented prior to the saccade, would be remapped to the left peripheral location, i.e. in the opposite direction of the saccade vector. Under the convergent remapping hypothesis, receptive fields are remapped towards the saccade target. Therefore, a test stimulus that is centrally presented would not be remapped to the left peripheral location, and no TAE is expected.

### Data Analysis

All data analyses were performed with MATLAB using the Palamedes Matlab toolbox for fitting psychophysical data (Prins and Kingdom, 2009). The significance threshold was set to 0.05. All data and code are available from Donders Institute for Brain, Cognition and Behavior Repository at https://data.donders.ru.nl/login/reviewer-30642337/C-C1x7OMV93tjHrM4l8A10sQaTm5NrqI-647Lqsj5As

### Outlier Criteria

#### Experiment 1

A total of 37,114 trials were obtained for experiment 1. Only successfully completed trials were considered in the further analyses. We excluded a trial from the analyses if a) fixation was broken before fixation displacement (7.75% of all trials), or b) the participant did not execute the required eye movements or missed the displaced fixation dot by more than 2° (10.87% of all trials). In the remaining trials, saccade latency was defined as the temporal distance between the onset of the fixation dot displacement and the initiation of the saccade that followed. Trials with saccade latencies shorter than 90 ms (0.23%) or longer than 500 ms (1.04%) were excluded. We also excluded trials whose response time was < 200 ms (0.3%) or more than 3 standard deviations above the subject’s mean response time (1.27%). Finally, trials in which the test stimulus was presented during the execution of the saccade were also excluded (15.71%). In total, 24,692 (81.55%) trials were included in the analysis.

#### Experiment 2

A total of 34,770 trials were obtained for Experiment 2. We excluded trials from further analyses if a) fixation was broken before fixation displacement (9.62%), or b) the participant did not execute the required eye movements or missed the displaced fixation dot by more than 2° (16.19%). Of the remaining trials, trials in which the saccade latency was < 90 ms (0.07%) or > 500 ms (1.04%) were excluded. We also excluded trials in which the button response time was < 200 ms (1.34%) or more than 3 standard deviations above the subject’s mean response time (1.08%). Finally, the trials in which test stimulus was presented during the saccade period were also excluded (16.75%). Together, 20,593 (79.83%) trials were included in the analysis.

#### Experiment 3

A total of 32,792 trials were obtained for Experiment 3. We excluded trials from the analyses if a) fixation was broken before fixation displacement (11.47% of all trials), or b) the participant did not execute the required eye movements or missed the displaced fixation dot by more than 2° (9.42% of all trials). Of the remaining trials, trials with saccade latencies shorter than 90 ms (0.07%) or longer than 500 ms (1.90%) were excluded. We also excluded trials whose response time was < 200 ms (5.32%) or more than 3 standard deviations higher than the subject’s mean response time (0.78%). Finally, trials in which test stimulus was presented during the execution of the saccade were also excluded (14.87%). In total, 20,820 (80.26%) trials were included in the analysis.

### Quantification of Time Bins

To plot the TAE magnitude as the function of time, we first separated all trials into two bins: one bin contained all trials in which the test stimulus was presented before saccade onset, whereas the other bin contained all trials in which the test stimulus was presented after saccade offset. Trials in which the test stimulus was presented during the saccade were removed. For both bins, the trials were then further subdivided into two time bins, respectively, with equal number of trials in each time bin. This resulted in a total of four time bins. In Experiment 1, mean test stimulus onset time with respect to saccade onset (for pre-saccadic trials) or saccade offset (for post-saccadic trials) was -133 ms (SD 73 ms), - 31 ms (SD 17 ms), 36 ms (SD 21 ms), and 115 ms (SD 28 ms) for each time bin, respectively. In Experiment 2, mean test stimulus onset time was -117 ms (SD 63 ms), -29 ms (SD 17 ms), 32 ms (SD 18 ms), and 104 ms (SD 26 ms) for each time bin, respectively. In Experiment 3, mean test stimulus onset time was -156 ms (SD 68 ms), -38 ms (SD 22 ms), 29 ms (SD 17 ms), and 97 ms (SD 26 ms) for each time bin, respectively.

### Quantification of Tilt Aftereffect

In order to quantify TAE magnitude, we fitted psychometric functions to the pooled group data. First, for each test stimulus location x adaptor tilt combination in each time bin, we expressed the proportion of “rightward” responses as a function of the test stimulus orientation with respect to vertical. For convenience, the leftward saccade trials were first collapsed with rightward saccade trials in each bin. Subsequently, we fitted cumulative normal distribution functions to this data. The point of subjective equality (PSE) was defined as the midpoint of the psychometric function, at which the test stimulus was perceived equally often as tilted to the right and left. The magnitude of TAE was then measured as half of the difference between the PSE of the leftward and rightward tilted adaptor conditions, for each time bin and each test stimulus location separately.

### Statistical Analyses

We used permutation tests to statistically compare: 1) differences of TAEs between time bins (before saccade), and 2) the interaction effect between locations (incorporating the initial fixation and the future saccade target location only) and the time bins (two time bins before eye movement) at the group level. First, to test for differences in TAEs between time bins, the condition labels of the first and second time bin of each participant were randomly shuffled. The resulting permutation group data was fitted with cumulative normal functions and was used to compute the difference in TAE between the time bins. This procedure was repeated 10,000 times. As p-values we report the proportion of permutations that led to an equal or more extreme TAE difference than the one we observed in the experiment. The exchangeability requirement for permutation tests is met, because under the null hypothesis of no difference in TAE between the first and second time bin, the condition labels are exchangeable. Second, in order to test for an interaction effect between locations and time bins, we first computed the differences of TAEs between initial fixation and saccade target location at each time bin, and then randomly shuffled the time bin labels of those differences for each participant. The exchangeability requirement for permutation tests is met, because under the null hypothesis of no interaction effect between locations and time bins, the TAE differences between locations should not be influenced by the time bin factor, and therefore the time bin labels are exchangeable. Again, this procedure was repeated for 10,000 times. As p-values we report the proportion of permutations that led to an equal or more extreme outcome than the one we observed in the experiment.

## Acknowledgements

T.H. was supported by a grant from China Scholarship Council (CSC, 201608330264). M.F. was supported by a grant from the European Union Horizon 2020 Program (ERC Starting Grant 678286, ‘‘Contextvision’’). F.P.d.L was supported by grants from The Netherlands Organization for Scientific Research (NWO; Vidi grant 452-13-016), the James S. McDonnell Foundation (Understanding Human Cognition, 220020373), and the EC Horizon 2020 Program (ERC starting grant 678286, ‘Contextvision’).

